# Target-specific machine-learning scoring functions boost structure-based virtual screening for *Pf*DHODH

**DOI:** 10.64898/2026.11.27.769741

**Authors:** Klaudia Caba, Pedro J. Ballester

## Abstract

*Plasmodium falciparum* dihydroorotate dehydrogenase (*Pf*DHODH) is a validated antimalarial target for which no inhibitor has yet been approved, and no target-specific machine learning scoring functions (ML SFs) have been reported for it. Here, we developed *Pf*DHODH-specific ML SFs for structure-based virtual screening from a small dataset of 73 experimentally confirmed actives and 86 inactives, supplemented with property-matched (PM) and property-unmatched (PU) decoys. We compared five algorithms, several protein-ligand interaction featurisation schemes and classification and regression formulations under two evaluation designs: a Butina clustering split and a UMAP-based split in which each cluster of actives was held out in turn. The best model, a random forest regressor trained on PLEC fingerprints combined with Morgan fingerprints, was the strongest under both designs, reaching a normalised enrichment factor at 1% (NEF1%) of 0.617 on the Butina test set, compared with 0.323 for Smina, the best-performing generic SF, and an average NEF1% of 0.656 across the UMAP clusters (Smina: 0.362). Regression generally outperformed classification, and deep learning models were unstable with the limited training data. Adding PU decoys to the training set substantially improved enrichment for most held-out clusters, whereas the cluster most dissimilar to the training actives remained poorly predicted. These results show that target-specific ML SFs can be developed for a target with scarce bioactivity data, provided that algorithms, featurisation and evaluation design are tested comprehensively.

## Introduction

Malaria remains a major global health burden, causing an estimated 610,000 deaths across 80 countries in 2024 alone, with the WHO African Region being the most affected and children under five accounting for 75% of deaths in the region [1]. Of the five parasite species responsible for malaria, *Plasmodium falciparum* is both the deadliest and the most prevalent across Africa. The continued treatment of malaria is complicated by widespread drug resistance, with the majority of antimalarial drugs developing resistance within approximately 20 years of their introduction [2]. *Pf*Dihydrofolate reductase (*Pf*DHFR), one of the most extensively studied antimalarial targets, is one of the examples of this pattern, having developed clinically significant resistance within two decades of the introduction of drugs targeting it. The RTS,S vaccine, while a notable advance, offers only modest efficacy (26 to 50%) and is not recommended for infants between 6 and 12 weeks of age [3], leaving a substantial gap in malaria prevention. Together, these limitations highlight the continued need for new antimalarial drug targets and inhibitors, ideally fast-acting, curative within three days and safe for use in pregnancy and early infancy [4].

Dihydroorotate dehydrogenase (DHODH) is an enzyme found in both humans (*Hs*DHODH) and the malaria-causing parasite *Plasmodium falciparum* (*Pf*DHODH). DHODH catalyses the conversion of dihydroorotate to orotate, the only redox reaction in de novo pyrimidine biosynthesis [5]. This reaction oxidises dihydroorotate to orotate while reducing the flavin cofactor FMN to FMNH_2_, which is subsequently reoxidised back to FMN by the co-substrate ubiquinone, itself reduced to ubiquinol (Figure 1). Inhibitors of DHODH act by preventing the binding of ubiquinone to FMNH_2_ [6].

**Figure 1:**
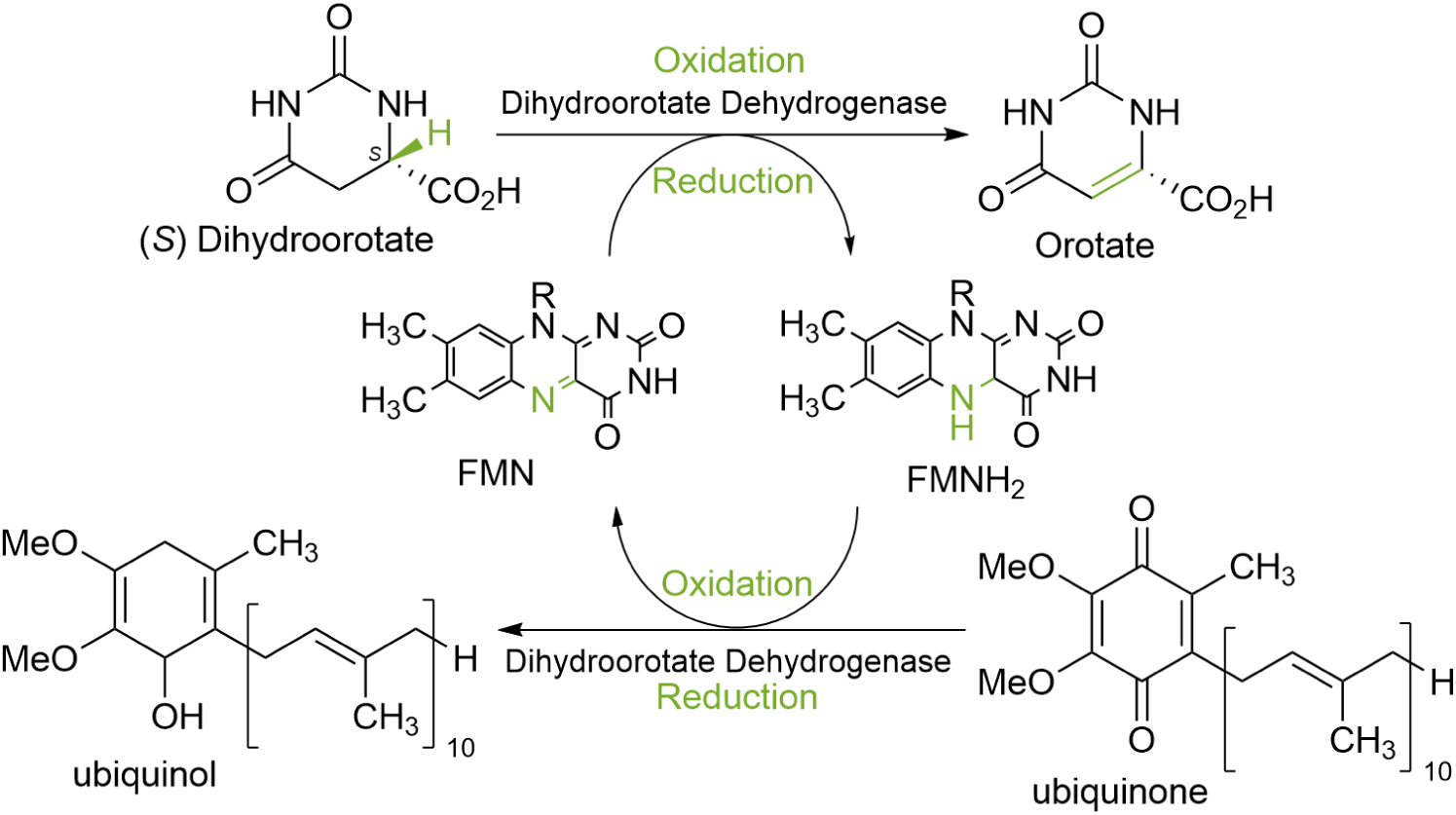
The redox reactions catalysed by dihydroorotate dehydrogenase (DHODH). Dihydroorotate is oxidised to orotate, the only redox reaction in de novo pyrimidine biosynthesis, while the flavin cofactor FMN is simultaneously reduced to FMNH_2_. FMNH_2_ is subsequently reoxidised to FMN by the co-substrate ubiquinone, which is itself reduced to ubiquinol. Adapted from [7].

*Pf*DHODH has emerged as a promising target for malaria treatment and prevention, since its inhibition blocks parasite growth at the blood stage by disrupting pyrimidine biosynthesis. While human cells can synthesise pyrimidines via both a salvage pathway and the de novo pathway, *P. falciparum* relies exclusively on de novo synthesis [8]. This absence of a salvage pathway renders the parasite particularly vulnerable to DHODH inhibition, as it has no alternative route to the nucleotide precursors required for RNA and DNA synthesis, cell growth and replication.

Despite the availability of alternate pyrimidine biosynthesis pathway in humans, selectivity between the human and parasite enzymes is essential. In humans, inhibiting de novo pyrimidine biosynthesis leads to immunosuppression, a desirable effect in the treatment of rheumatoid arthritis and in organ transplantation, but a dangerous one in the context of malaria infection, where a functioning immune response is needed to clear the parasite [9]. Achieving this selectivity is possible because *Pf*DHODH can be inhibited at either of two sites: the dihydroorotate (DHO) substrate-binding site or the ubiquinone (CoQ) co-substrate-binding site. The DHO site is highly conserved across species, making it a poor target for species-selective inhibition, whereas the CoQ site shows considerable structural variability between *Hs*DHODH and *Pf*DHODH, making it the preferred target for the development of *Pf*DHODH-selective inhibitors. The CoQ-binding tunnel can be divided into three regions, the mouth, waist and end, and structural and chemical differences across these regions underlie the species selectivity achievable at this site [10]. The mouth of *Hs*DHODH, for example, is notably broader than that of *Pf*DHODH, a difference attributed to a slight structural kink introduced by Leu42 in the human enzyme. The specific non-conserved residues contributing to these structural differences across the tunnel are shown in Figure 2A. Five of them (Leu172, Leu176, Phe188, Ile263 and Ile272) form hydrophobic contacts with DSM265, a selective *Pf*DHODH inhibitor, whereas its hydrogen bonds involve His185 and Arg265 (Figure 2B). Differences at these positions are therefore likely to contribute to inhibitor selectivity.

**Figure 2:**
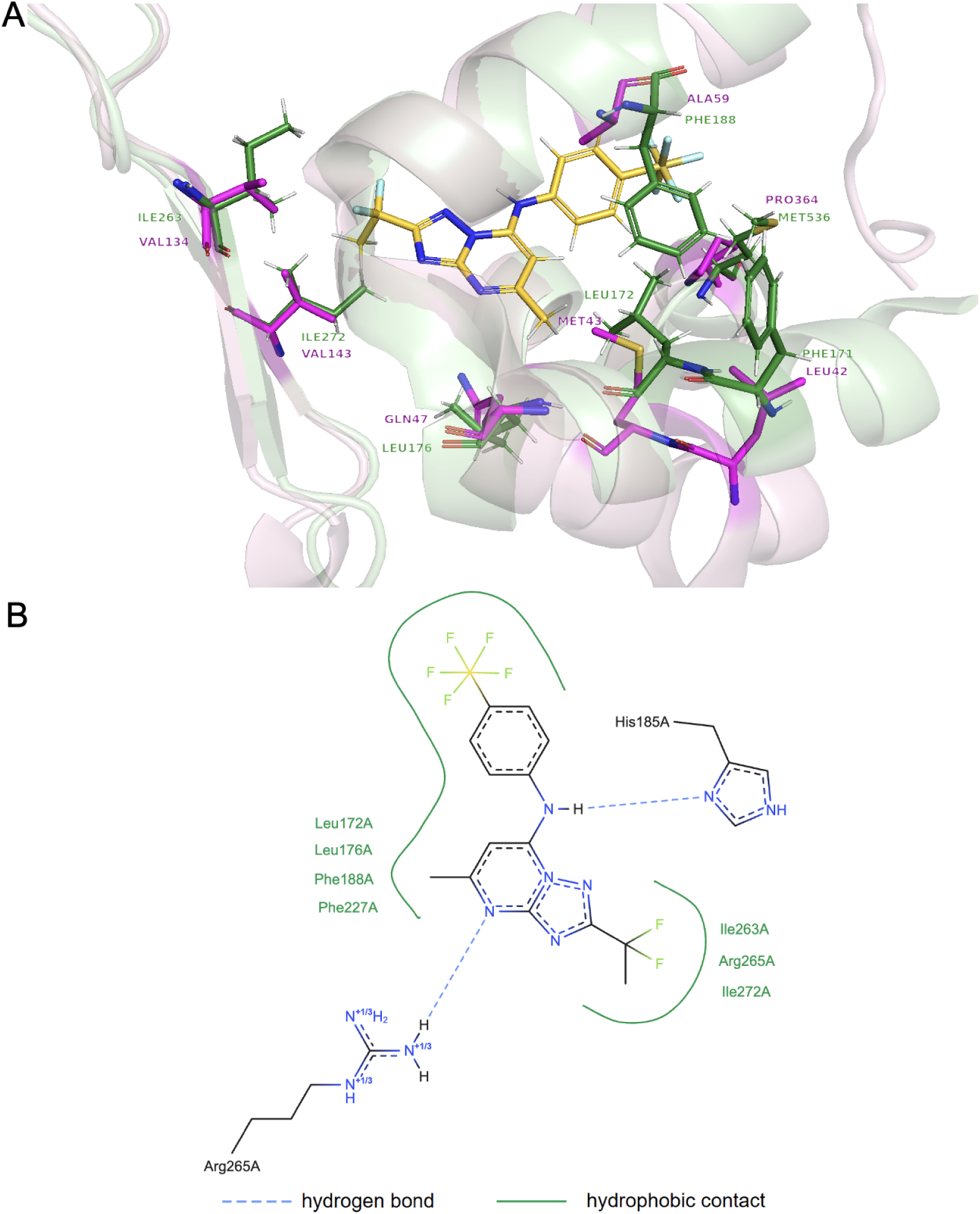
(A) Superposed CoQ-binding tunnels of *Pf*DHODH (green; PDB ID 4RX0) and *Hs*DHODH (magenta; PDB ID 2FPY), showing the non-conserved residues as sticks: Phe171/Leu42, Met536/Pro364, Leu172/Met43, Phe188/Ala59, Leu176/Gln47, Ile263/Val134 and Ile272/Val143 (*Pf*DHODH/*Hs*DHODH). The *Pf*DHODH inhibitor DSM265 is shown in gold. (B) 2D diagram of the interactions between DSM265 and *Pf*DHODH. Interaction types are indicated in the legend.

Numerous chemical classes of *Pf*DHODH inhibitors have been reported over the past two decades, spanning triazolopyrimidines, N-phenylbenzamides, dihydrothiophenones, pyrimidones and several others [11]. Baldwin et al. performed the first enzyme-based high-throughput screen to identify *Pf*DHODH inhibitors, from which a triazolopyrimidine-based compound, DSM265, emerged as the most advanced candidate, progressing to phase IIa clinical trials [12]. Its development was however halted following safety signals of teratogenicity and testicular toxicity [13]. Despite this progress, no *Pf*DHODH inhibitor has yet reached clinical approval, underscoring the continued need for the discovery of novel, selective inhibitors with improved safety profiles.

Several computational studies have focused on identifying novel *Pf*DHODH inhibitors, most employing ligand-based QSAR or pharmacophore modelling alongside molecular docking against available *Pf*DHODH crystal structures. Hou et al. developed QSAR models using SVM and multilinear regression trained on a dataset of 255 *Pf*DHODH inhibitors [14]. Pavadai et al. combined pharmacophore-based virtual screening with molecular docking to identify NSC336047, a selective *Pf*DHODH inhibitor with negligible activity against *Hs*DHODH (IC_50_ *>* 100 *µ*M) and an IC_50_ of 26 *µ*M against parasite growth [15]. More recently, Thomas et al. introduced RL-MSVP, a pipeline combining reinforcement learning-based molecular generation with molecular docking and molecular dynamics simulations, identifying 50 novel azole-fused pyrimidine *Pf*DHODH inhibitors with predicted binding affinities exceeding that of DSM265 [16]. Despite this body of work, no generic or target-specific ML SF has previously been developed or applied to *Pf*DHODH in the peer-reviewed literature, motivating the scope of the work presented here.

## Methodology

### Data collection and preprocessing

Bioactivity data for *Pf*DHODH (UniProt ID: Q08210) was collected from PubChem and ChEMBL, including measurements reported as IC_50_, EC_50_ and K*_i_*. Only compounds associated with the Inhibitor modulator type were retained. Data was curated according to the following steps. Firstly, molecules with missing activity concentrations or no SMILES were removed. Any entries associated with anything other than ‘Confirmatory’ in the assay type were also deleted. Moreover, any compounds which would fall under the active category but with relationship type reported as “*>*” were discarded since the true concentration is potentially greater than the one reported introducing uncertainty as to whether the compound is truly active. Compounds for which entries suggested contradictory activity classifications were discarded. Additionally, any compounds present across multiple entries but with ambiguous information were removed. After data cleaning steps, activity concentrations were log transformed to pIC_50_ values.

An activity threshold of 1 *µ*M (pIC_50_ of 6) was applied to label actives and inactives yielding 73 experimental actives and 89 experimental inactives. The threshold of 1 *µ*M is a regularly used value in this field [17, 18, 19, 20]. Three of these inactives were subsequently removed because EHIGN graphs could not be generated for them, leaving 86 experimental inactives.

The 4RX0 crystal structure of *Pf*DHODH (resolution 2.25 Å) was selected owing to its relatively high resolution, the absence of mutations and the presence of the co-crystallised inhibitor DSM265. Following selection, protein structure was prepared for docking using the Dock Prep tool in UCSF Chimera (v1.17) [21]. All default Dock Prep options were enabled with the exception of “Delete solvent”, “Delete non-complexed ions” and “Add charges”, as recommended by dos Santos et al. [22]. Along with the prepared protein template, the bound ligand DSM265 was extracted from the co-crystallised structure to guide the binding site in molecular docking experiments.

Property-matched (PM) decoys were generated for all 73 actives using DeepCoy, giving 3650 decoys, of which 194 were later removed because of the EHIGN graph generation failures, leaving 3456. As experimentally confirmed inactives were scarce, 1500 property-unmatched (PU) decoys were additionally collected from the ZINC 22 database [23] via the CarteBlanche interface to increase the structural diversity of the inactive class available for model training. The way PM and PU decoys were allocated to training and test sets depends on the data splitting strategy and is described in the corresponding Butina and UMAP sections. The final dataset therefore comprised 73 experimental actives, 86 experimental inactives, 3456 PM decoys and 1500 PU decoys, with the PU decoys used only in a subset of the UMAP-based experiments.

Ligand SMILES were protonated at pH 7.4 using OpenBabel [24], and 3D coordinates were then generated with the RDKit functions `rdDistGeom.ETKDGv3` and `rdDistGeom.EmbedMolecule` [25]. Molecular docking was performed with Smina using the prepared 4RX0 protein template, with DSM265 defining the binding site. A cubic search space of 30 Å along each axis was defined to provide sufficient space for ligand sampling. Only the top-ranked pose per ligand, according to the Smina docking score, was retained. The default value of 8 was used for `--exhaustiveness`. A random seed (`--seed`) of 0 was set for all docking experiments to ensure reproducibility.

### Featurisation

Six featurisation schemes were investigated. The first group is based on PLEC - protein-ligand extended connectivity fingerprints [26]: PLEC alone, PLEC combined with Smina terms (Smina-PLEC), and PLEC combined with ligand-only features, either physicochemical descriptors (physchem-PLEC) or Morgan fingerprints (MF-PLEC). The second group comprises two ECIF-based schemes, multi-shelled and weighted ECIF. PLEC fingerprints were extracted using the Open Drug Discovery Toolkit (ODDT) [27] package using the (`depth_ligand`) of 1 and (`depth_protein`) of 5. Smina terms (`gauss(o=0,_w=0.5,_c=8`), `gauss(o=3,_w=2,_c=8`), `repulsion(o=0,_c=8)`, hydrophobic interactions between atoms `hydrophobic(g=0.5,_b=1. 5,_c=8)`, hydrogen bonding `non_dir_h_bond(g=-0.7,_b=0,_c=8)` and number of torsional angles `num_tors_div`) were extracted from Smina-docked protein-ligand complexes. Multi-shelled and weighted ECIF features were computed using the original implementation provided by the authors [28]. Morgan fingerprints (512 bits, radius 2) were computed using `rdFingerprintGenerator.GetMorganGenerator` function in rdkit. Physicochemical descriptors were computed by calculating all descriptors available in the `rdMolDescriptors.Properties` package with the exception of two descriptors relating to stereocentres, following the approach used by the authors of a generic ML SF vScreenML2 [29].

In the final stage of the UMAP experiments, recursive feature elimination (RFE) was applied to the PLEC features (4092 in total), retaining the 500 highest-ranked features. Selection was performed separately for each of the four UMAP splits, using only the training set of that split (including the PU decoys), and the final RF, XGB and SVM models were trained on the selected features, with hyperparameters optimised as before. The number of retained features was fixed a priori at 500, approximately 12% of the original set and of a similar size to the 512-bit Morgan fingerprints used in the MF-PLEC scheme. It was not optimised, so the results should be taken as an indication of the effect of feature selection and not as the best achievable with it. Features were removed one at a time, the sklearn default, so that the importances were recomputed after each elimination.

### Data splitting strategies

Owing to the limited amount of collected data, two clustering-based data splitting strategies, Butina and UMAP, were used for model training and evaluation. They were not applied simultaneously: the UMAP-based approach was developed after drawbacks of the Butina-based split were identified, as discussed in the Butina results sections. Within each strategy, all target-specific models were trained on the same training set, and all methods, including the off-the-shelf reference SFs that require no training, were evaluated on the same test set(s).

#### Butina-clustering-based split

In this first approach, a single training-test split was formed with the aim of testing the generalisation of the models to structurally diverse actives. The 73 experimental actives were clustered with the Butina algorithm using Morgan fingerprints (2048 bits, radius 2) and a Tanimoto similarity threshold of 0.6. This produced 15 clusters, the largest of which contained 33 molecules (Table 1). The actives in the largest cluster formed the training actives (33), and those in the remaining clusters formed the test actives (40). All 86 experimental inactives were assigned to the training set, giving a training set of 119 compounds (33 actives and 86 inactives). The test set comprised the 40 test actives and all 3456 PM decoys. As PM decoys were generated for all 73 actives, these include decoys matched to the training actives. The active fraction in the test set is therefore 1.1% (approximately 86 decoys per active), higher than the 50 decoys per active used in decoy generation. PU decoys were not used in this strategy.

**Table 1:** Butina clusters of the 73 experimental actives (Morgan fingerprints, Tanimoto similarity threshold of 0.6).

| **Cluster(s)** | **Actives per cluster** | **Set** |
|---|---|---|
| 0 | 33 | Training |
| 1 | 8 | Test |
| 4, 5 | 7 each | Test |
| 3, 8, 9 | 3 each | Test |
| 7 | 2 | Test |
| 2, 6, 10, 11, 12, 13, 14 | 1 each | Test |

#### UMAP clustering split

In the second strategy, the 73 experimental actives were clustered in a UMAP embedding of their Morgan fingerprints (2048 bits, radius 2). As UMAP does not fix the number of clusters, the number of neighbours (5, 10, 15 or 20) and the number of clusters (2 to 7) were optimised jointly, giving 24 combinations, using default values for all other UMAP parameters. The combination with the best trade-off between cluster separation and compactness was 5 neighbours and 4 clusters, which gave clusters of 24, 17, 25 and 7 actives (clusters 0 to 3), a silhouette score of 0.859, a Calinski-Harabasz score of 1950.5 and a Davies-Bouldin score of 0.176.

Instead of a single training-test split, four splits were formed, each holding out one cluster as the test actives (Figure 3). The training set of each split comprised the actives from the remaining three clusters and all 86 experimental inactives. In a subset of experiments, the 1500 PU decoys were added to the training set to increase the diversity of the inactive class. Two test sets were built for each split, both containing the held-out actives. The all-decoys test set contained all 3456 PM decoys, including those matched to the training actives, which provides a larger and more chemically diverse set of decoys that is identical across splits. The matched-decoys test set contained only the PM decoys matched to the held-out actives (1161, 758, 1206 and 331 decoys for clusters 0 to 3), which is the more common design for retrospective benchmarks. The active fractions are 0.69, 0.49, 0.72 and 0.20% in the all-decoys test sets and 2.0 to 2.2% in the matched-decoys test sets. Experimental inactives and PU decoys were never included in test sets.

**Figure 3:**
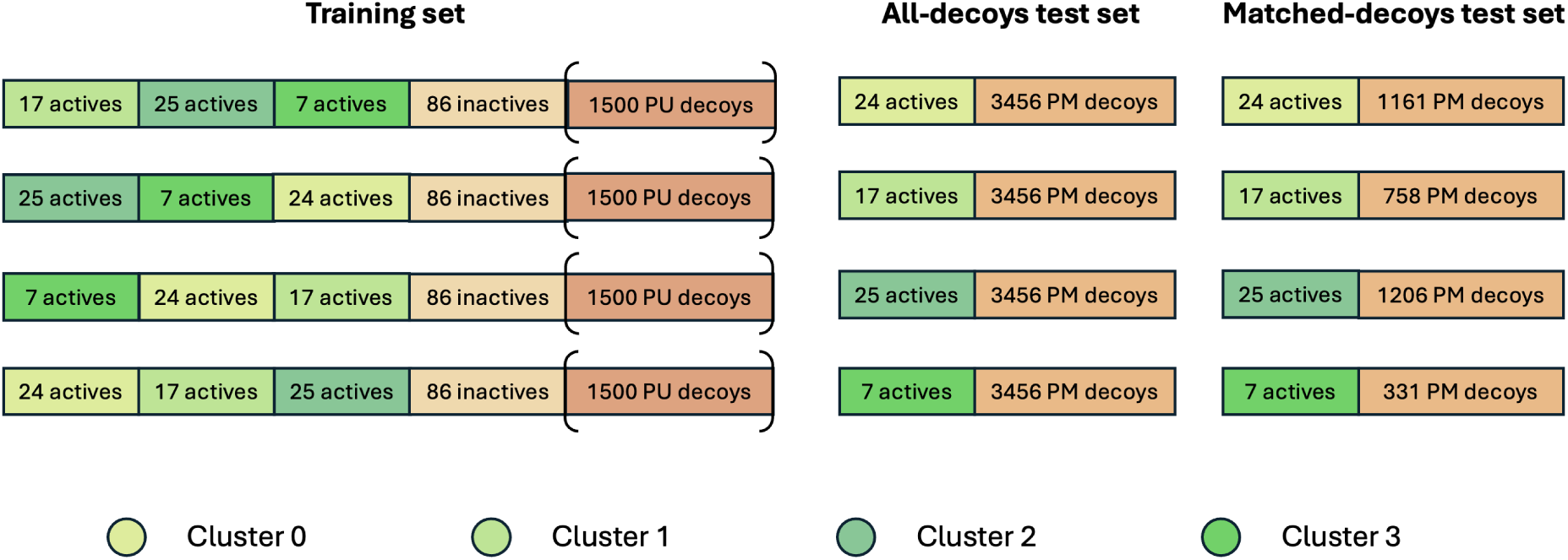
Training and test sets for the four UMAP-based splits. In each split, one cluster of actives (clusters 0 to 3) is held out as the test actives, and the training set comprises the actives of the other three clusters and the 86 experimental inactives. The 1500 PU decoys were added to the training set in a subset of experiments. The all-decoys test set contains the held-out actives and all 3456 PM decoys, and the matched-decoys test set contains the held-out actives and only the PM decoys matched to them.

### Model training

Five ML algorithms were trained with the Butina clustering split: RF, XGB, SVM, ANN and DNN. For each algorithm, both classification and regression variants were trained and evaluated. With the UMAP clustering split, only RF, XGB and SVM regression models were used, a choice motivated by the Butina results. Models were implemented with the Python packages (`sklearn` [30], `tensorflow`), and each was trained 10 times with different random seeds, with the median performance reported. Hyperparameters were optimised as described in the next section. The training and test sets, featurisation schemes and other design choices for the two splits are compared in Table 2.

**Table 2:**
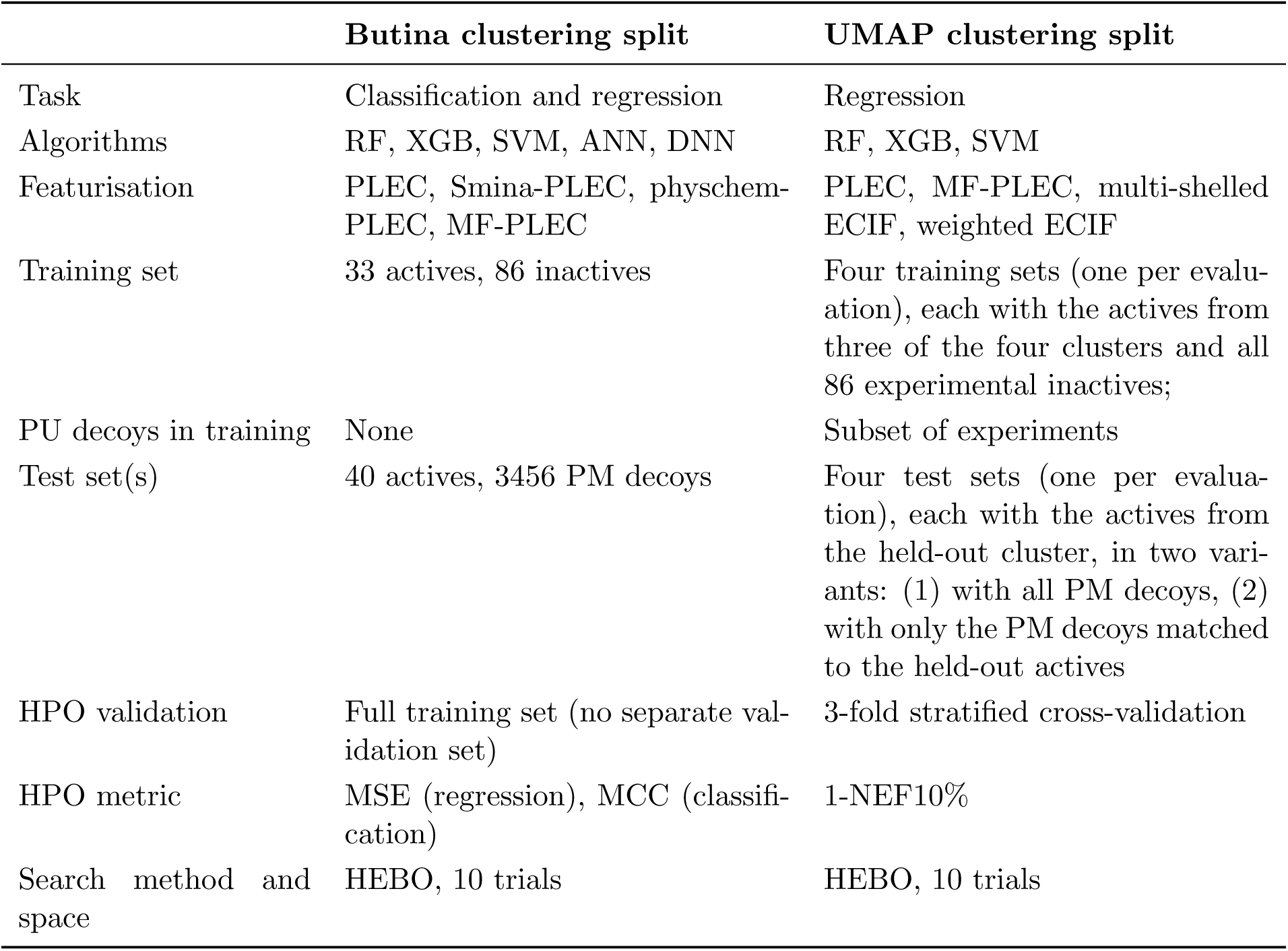
Comparison of the model development set-up for the Butina and UMAP clustering splits.

|  | **Butina clustering split** | **UMAP clustering split** |
|---|---|---|
| Task | Classification and regression | Regression |
| Algorithms | RF, XGB, SVM, ANN, DNN | RF, XGB, SVM |
| Featurisation | PLEC, Smina-PLEC, physchem-PLEC, MF-PLEC | PLEC, MF-PLEC, multi-shelled ECIF, weighted ECIF |
| Training set | 33 actives, 86 inactives | Four training sets (one per evalu-ation), each with the actives from three of the four clusters and all 86 experimental inactives; |
| PU decoys in training | None | Subset of experiments |
| Test set(s) | 40 actives, 3456 PM decoys | Four test sets (one per evalua-tion), each with the actives from the held-out cluster, in two vari-ants: (1) with all PM decoys, (2) with only the PM decoys matched to the held-out actives |
| HPO validation | Full training set (no separate val-idation set) | 3-fold stratified cross-validation |
| HPO metric | MSE (regression), MCC (classifi-cation) | 1-NEF10% |
| Search method and space | HEBO, 10 trials | HEBO, 10 trials |

### Hyperparameter optimisation

Hyperparameters were optimised with HEBO [31] using 10 trials per model. The search spaces are given in Table 3.

**Table 3:**
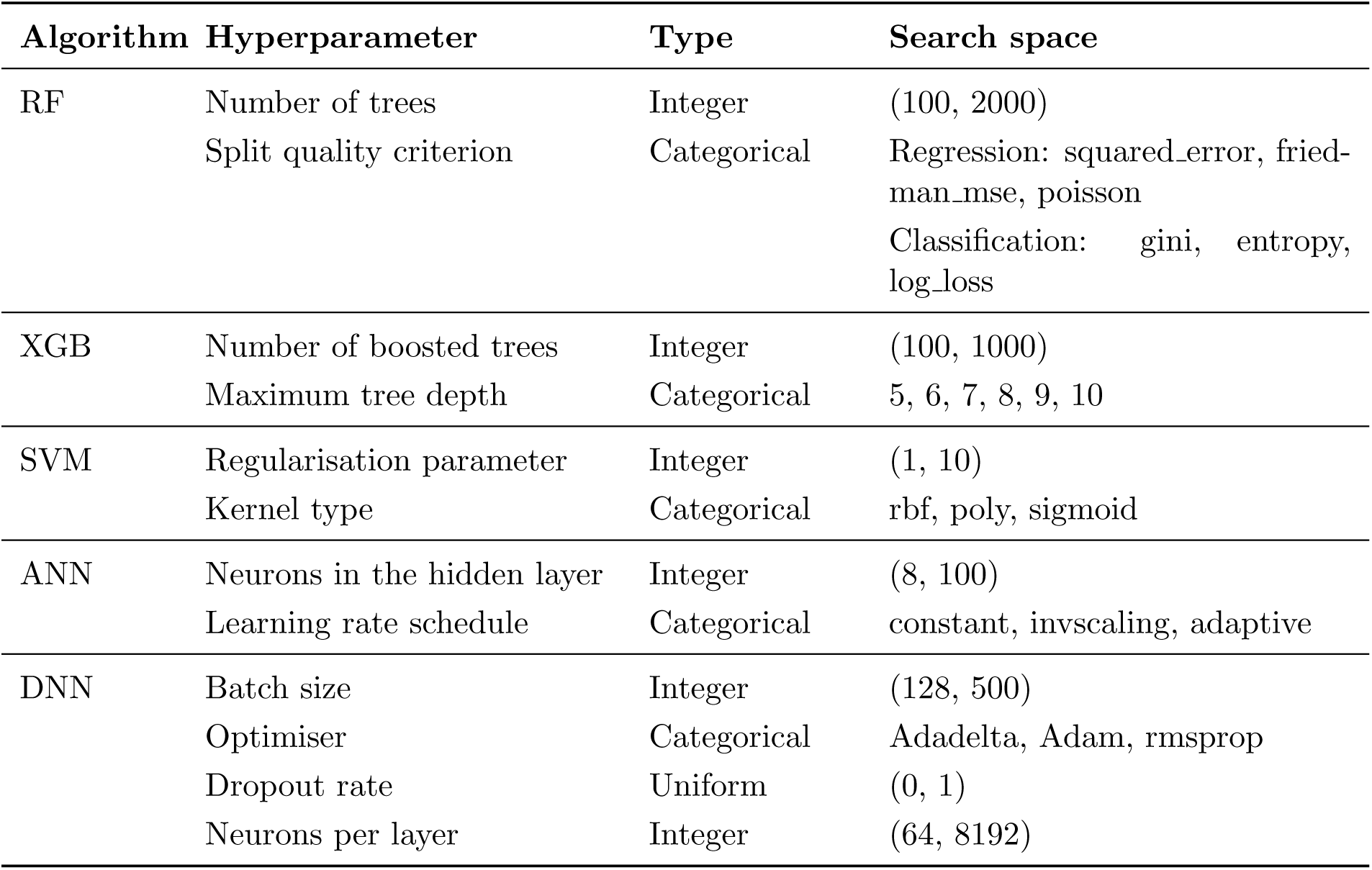
Hyperparameters and search spaces optimised with HEBO. The same spaces were used for the Butina and UMAP clustering splits. The ANN and DNN spaces as well as RF classification criteria apply to the Butina split only.

| **Algorithm** | **Hyperparameter** | **Type** | **Search space** |
|---|---|---|---|
| RF | Number of trees | Integer | (100, 2000) |
|  | Split quality criterion | Categorical | Regression: squared_error, fried-man_mse, poisson<br>Classification: gini, entropy, log_loss |
| XGB | Number of boosted trees | Integer | (100, 1000) |
|  | Maximum tree depth | Categorical | 5, 6, 7, 8, 9, 10 |
| SVM | Regularisation parameter | Integer | (1, 10) |
|  | Kernel type | Categorical | rbf, poly, sigmoid |
| ANN | Neurons in the hidden layer | Integer | (8, 100) |
|  | Learning rate schedule | Categorical | constant, invscaling, adaptive |
| DNN | Batch size | Integer | (128, 500) |
|  | Optimiser | Categorical | Adadelta, Adam, rmsprop |
|  | Dropout rate | Uniform | (0, 1) |
|  | Neurons per layer | Integer | (64, 8192) |

#### Butina clustering split

All five algorithms (RF, XGB, SVM, ANN and DNN) were optimised, in both regression and classification variants. The objective was the MSE for regression models, and the MCC for classification models. As the training set was limited in size to hold out a separate validation set for hyperparameter tuning, hyperparameters were tuned on the full training set.

#### UMAP clustering split

Only the RF, XGB and SVM regression models were used, so the search spaces for these three algorithms in Table 3 apply. Hyperparameters were optimised within each training set by stratified 3-fold cross-validation (`StratifiedKFold` from `sklearn`, with shuffling and a fixed random seed of 42). The folds were stratified by active or inactive label, so that the proportion of actives is preserved in each fold and both classes are present in every training and validation partition, which matters given the small size of the training sets. In each iteration, models were trained on two folds and evaluated on the remaining one. 1-NEF10% was used as the objective function to be minimised. NEF10% was chosen over NEF1% because the larger cut-off gives a less coarse metric on small validation folds.

### Reference scoring functions

The target-specific ML SFs were compared with three reference SFs: Smina, gnina and EHIGN. Smina, the classical SF used for docking, served as the baseline and was applied in both Butina and UMAP experiments, with compounds ranked by their Smina docking scores. Gnina, a generic ML SF, was also applied throughout to rescore the Smina-docked poses, and both its outputs, CNNscore and CNNaffinity, were used for ranking.

EHIGN was applied with the Butina clustering split only. As no pretrained generic EHIGN model for SBVS was publicly available, it was trained on the *Pf*DHODH training set, and the Butina test set was used for evaluation. EHIGN was not carried forward to the UMAP clustering split because it performed poorly with the Butina split, similarly to the ANN and DNN models: DL architectures are data-hungry and are unlikely to perform well on the small training sets available for this target.

For EHIGN, interaction graphs were generated from the docked complexes using the code provided by the original authors [32]. Graph generation failed for 197 ligands (3 experimental inactives and 194 PM decoys). To ensure that all methods were trained and evaluated on identical compounds, these ligands were removed from the dataset.

### Evaluation

Performance was evaluated using the enrichment factor at 1% (EF1%) and its normalised version (NEF1%). NEF1% was used as the main metric, because it accounts for the different numbers of actives and decoys in the test sets and so allows comparison between them. All methods were evaluated on the same test sets: the Butina test set, and for the UMAP clustering split the all-decoys and matched-decoys test sets of each of the four splits. Compounds were ranked by the Smina docking score for Smina, by CNNscore and CNNaffinity for gnina, by the predicted activity for the regression models, and by the predicted probability of being active for the classification models and EHIGN. For the ML models trained with random seeds, the median over 10 runs is reported. For the UMAP clustering split, results are reported for each split and averaged across the four splits.

## Results and Discussion

### Butina approach

Butina clustering is widely used to split datasets for model evaluation and it was the first strategy applied here. The training set comprised 33 actives from the largest cluster and 86 experimental inactives, and the test set comprised the 40 remaining actives and 3456 PM decoys (1.1% actives). Twenty regression and twenty classification models (five algorithms, each with four featurisation schemes) were trained and evaluated using the median NEF1% over 10 runs. This section reports the results from this approach and then examines its limitations that motivated the UMAP-based approach.

### Model performance across featurisation schemes

Performance varied more with the featurisation scheme than with the task (Figure 4). PLEC alone gave a reasonable baseline for most algorithms, with the strongest results for RF and DNN regressors (NEF1% of 0.588 and 0.559) and the ANN classifier (0.500).

**Figure 4:**
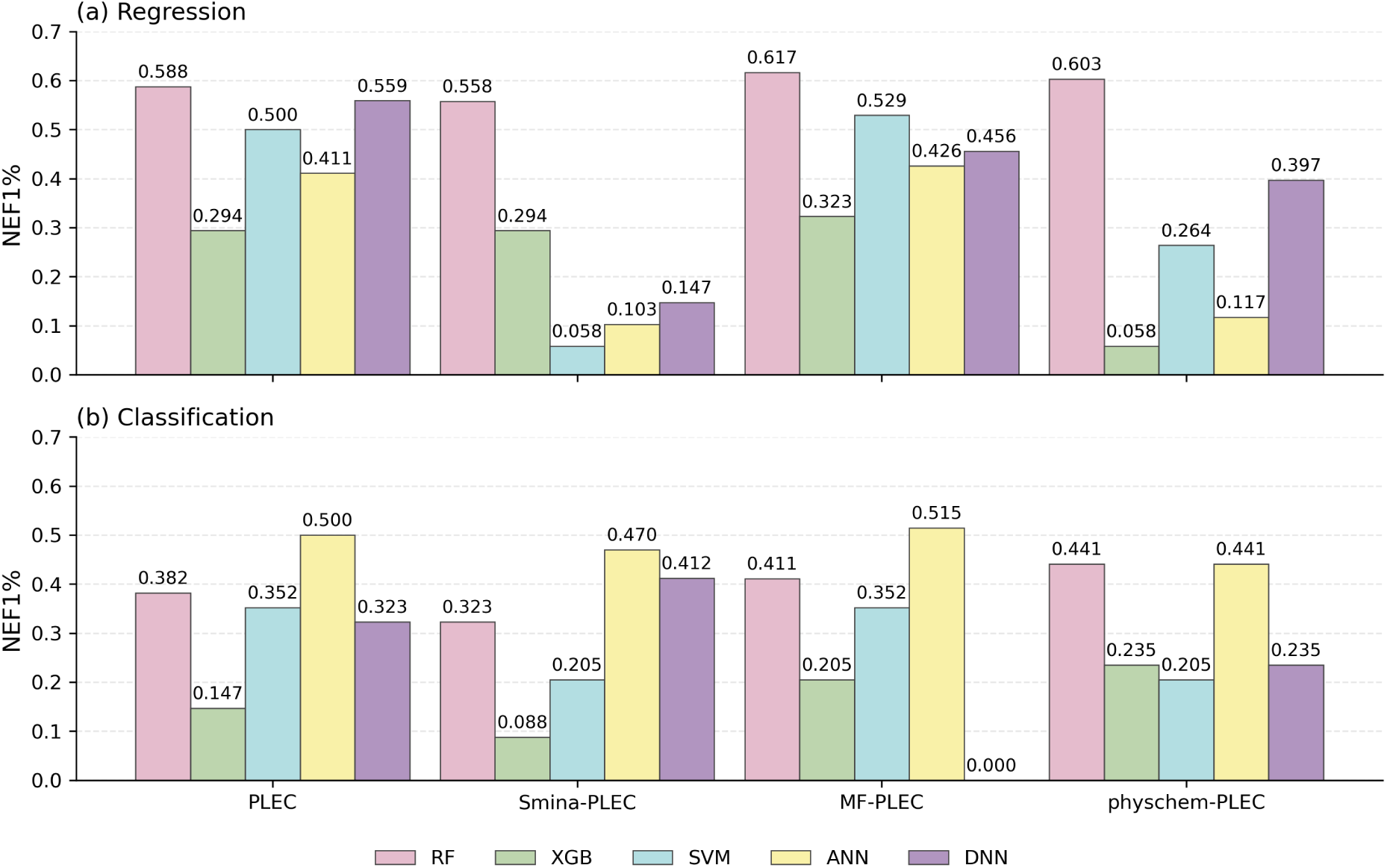
Median NEF1% over 10 runs for the *Pf*DHODH-specific (a) regression and (b) classification models on the Butina clustering split test set (40 actives and 3456 PM decoys), across algorithms and featurisation schemes.

Adding Smina terms to PLEC mostly lowered performance. For regression, NEF1% fell for RF, SVM, ANN and DNN and was unchanged for XGB, with the largest drops for SVM (0.500 to 0.058), ANN (0.411 to 0.103) and DNN (0.559 to 0.147). For classification, performance fell for RF, XGB, SVM and ANN, and only the DNN classifier improved (0.323 to 0.411). One possible explanation is the difference in feature scale: PLEC features are integer counts with a maximum of 35 in this dataset, whereas Smina terms reach values of up to 2306.305. Scale differences affect SVM, ANN and DNN but not tree-based models. Normalising the Smina terms might therefore recover some of the lost performance for these algorithms, but this was beyond the scope of this work.

Adding Morgan fingerprints to PLEC gave the most consistent improvement. For regression, NEF1% increased for RF, XGB, SVM and ANN, with only DNN decreasing (0.559 to 0.456), and MF-PLEC gave the highest value for four of the five algorithms. For classification, it improved RF, XGB and ANN, left SVM unchanged (0.352) and reduced the DNN value to 0.

Adding physicochemical descriptors had mixed effects. For regression, it improved only RF (0.588 to 0.603) and lowered performance for XGB, SVM, ANN and DNN, in some cases substantially (XGB from 0.294 to 0.058). For classification, it gave the highest NEF1% for RF (0.441) and XGB (0.235), but lowered performance for SVM, ANN and DNN.

Overall, RF was the most robust algorithm across schemes, with regression NEF1% between 0.558 and 0.617. XGB, SM, ANN and DNN were far more sensitive to the choice of features.

### Classification versus regression

Regression models generally outperformed classification models. The best regression model (RF with MF-PLEC, NEF1% = 0.617) exceeded the best classification model (ANN with MF-PLEC, 0.515), and for four of the five algorithms the best regression variant was better than the best classification variant: RF (0.617 versus 0.441), XGB (0.323 versus 0.235), SVM (0.529 versus 0.352) and DNN (0.559 versus 0.412). ANN was the exception (0.426 for regression versus 0.515 for classification). A plausible explanation is that regression uses the measured activity values and so retains more information than the binary labels obtained by applying the 1 *µ*M threshold, which is particularly relevant with only 33 training actives.

The classification models also showed a failure mode that regression did not. The median NEF1% of the DNN classifier with MF-PLEC fell to 0. Inspection of the ranked test set showed that the model assigned a probability of exactly 1 to a large fraction of compounds. Compounds with identical probabilities were ordered arbitrarily, and the top 34 compounds contained only decoys. Probability outputs saturate more easily than continuous predictions, which makes ranking by classification probability fragile when many compounds tie.

### Model performance across algorithms

The five algorithms differed in both the level and the stability of their performance. RF was the strongest and most robust. Across the four featurisation schemes, the RF regressor reached NEF1% values of 0.558 to 0.617 (Figure 4), and across 10 runs with different random seeds its NEF1% with PLEC varied only between 0.558 and 0.588, a difference of a single active in the top 1% of the ranked test set. XGB and SVM were also stable across seeds, but they were more sensitive to the choice of features: the XGB regressor ranged from 0.058 to 0.323 across schemes, and the SVM regressor fell from 0.500 to 0.058 when Smina terms were added to PLEC.

The ANN and DNN regression models were the least stable, in two respects. First, their performance changed sharply with the featurisation scheme. The DNN regressor ranged from 0.147 to 0.559 and the ANN regressor from 0.103 to 0.426. Second, their performance varied considerably between random seeds even with a fixed featurisation. For example, with PLEC, the NEF1% of the ANN regressor ranged from 0.294 to 0.529 across 10 runs and that of the DNN regressor from 0.352 to 0.617 (Figure 5). This is a spread of several actives in the top 1%, whereas RF, XGB and SVM varied by at most one active. Seed-to-seed variability was examined for PLEC regression models only, as the baseline featurisation common to all models.

**Figure 5:**
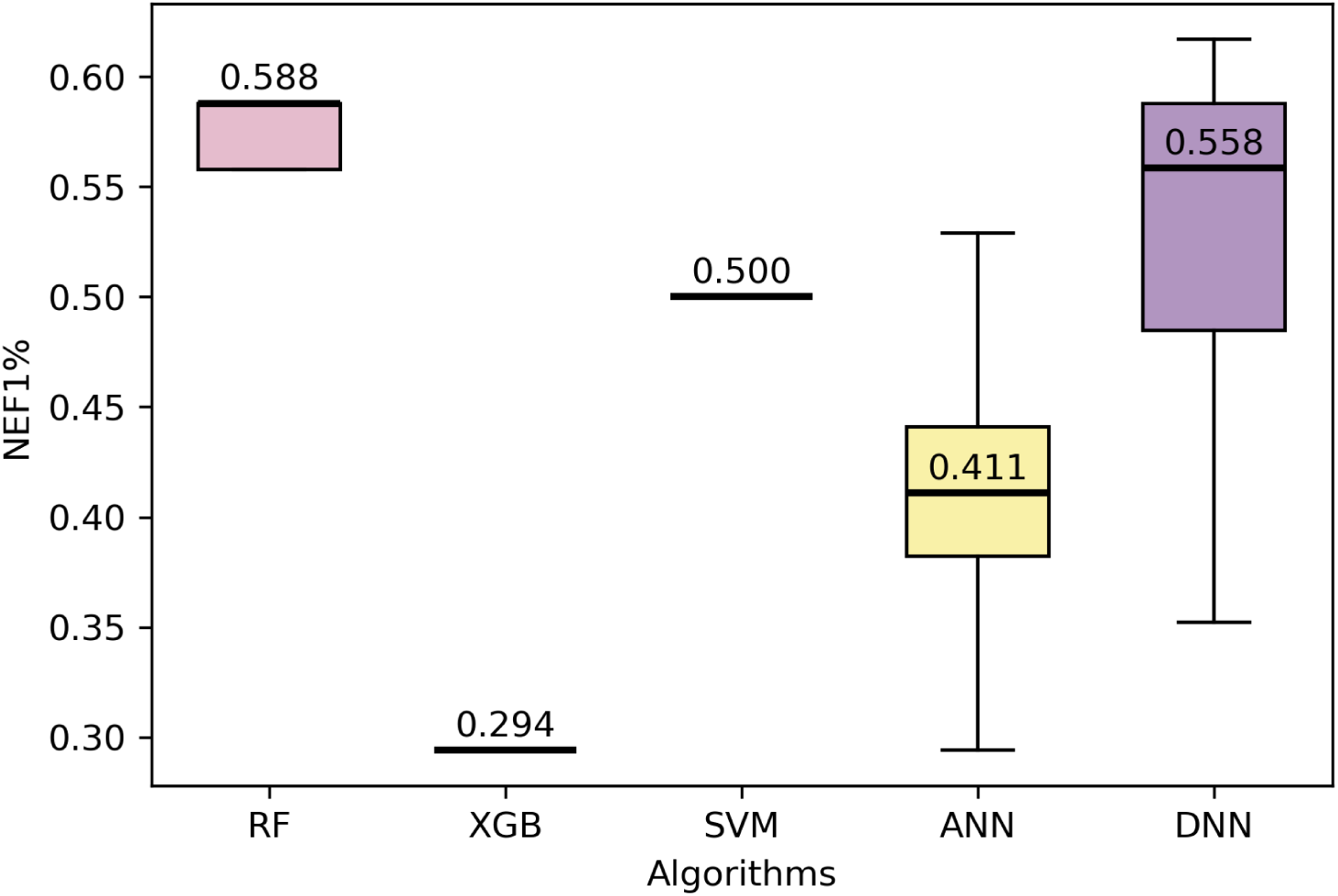
NEF1% of the regression models trained on PLEC features across 10 runs with different random seeds on the Butina clustering split test set. The line marks the median, the box the interquartile range and the whiskers the minimum and maximum. XGB and SVM gave identical values in all runs, and RF varied by a single active in the top 1% (0.558 to 0.588).

This instability means that a single run of an ANN or DNN regressor could give a misleadingly high or low result, and even the median over 10 runs conceals a wide range of outcomes. With a training set of only 119 compounds (33 actives), this variability is consistent with insufficient data for these models. Because the variability arises on identical training data, a consensus model that returns, for each test compound, the median of the predictions of the 10 models could give better or at least more stable DNN performance, and is worth testing in future work. Among the regression models, RF, XGB and SVM were stable across seeds, and RF achieved the highest performance, whereas the ANN and DNN regressors were sensitive to the random seed.

### Comparison to reference scoring functions

Among the reference SFs, Smina performed best (NEF1% = 0.323), followed by EHIGN (0.294), CNNaffinity (0.147) and CNNscore (0.029). When comparing their performance to the ten target-specific ML SFs with PLEC featurisation strategy as baseline, seven models outperformed Smina, PLEC-DNN classifier matched it (NEF1% = 0.323) and both XGB regressor and classifier performed worse (NEF1% = 0.294 and NEF1% = 0.147, respectively) - Table 4 and Figure 6. Eight of the ten models outperformed EHIGN. The best model was the RF regressor with MF-PLEC (EF1% = 52.5, NEF1% = 0.617), which recovers approximately 21 of the 40 actives within the top 34 compounds, almost twice the NEF1% of Smina. EHIGN did not outperform Smina, and its result is below that of most of the simpler models, consistent with the small training set limiting a more complex architecture.

**Figure 6:**
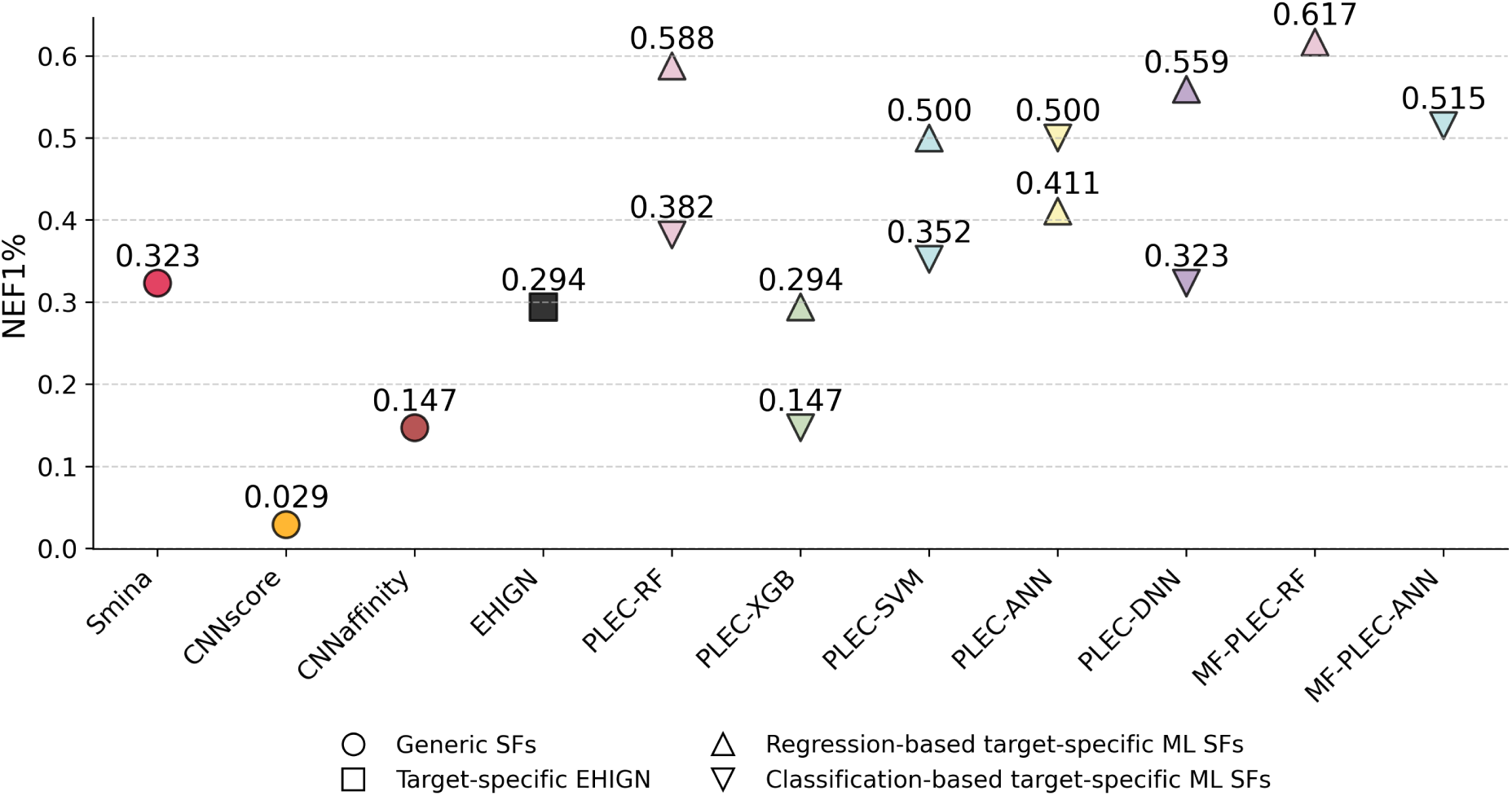
NEF1% of the reference SFs and the *Pf*DHODH-specific ML SFs on the Butina clustering split test set (40 actives and 3456 PM decoys). Each algorithm is shown trained with PLEC features, together with the best-performing regression (RF) and classification (ANN) models overall, both trained with MF-PLEC. Values for target-specific models are medians over 10 runs, except for EHIGN, which is a single model.

**Table 4:** Performance of the reference SFs and the *Pf*DHODH-specific ML SFs on the Butina clustering split test set (40 actives and 3456 PM decoys). Target-specific models are shown for each algorithm and task using PLEC as the common featurisation, together with the best-performing regression (RF) and classification (ANN) models overall, both trained with MF-PLEC. Values for target-specific models are medians over 10 runs, except for EHIGN, which is a single model.

| **Scoring function** | **Type** | **EF1%** | **NEF1%** |
|---|---|---|---|
| Smina | Regression | 27.5 | 0.323 |
| CNNscore | Classification/Regression | 2.50 | 0.029 |
| CNNaffinity | Classification/Regression | 12.5 | 0.147 |
| EHIGN (*Pf*DHODH-specific) | Classification | 25.0 | 0.294 |
| PLEC-RF | Regression | 50 | 0.588 |
| PLEC-XGB | Regression | 25 | 0.294 |
| PLEC-SVM | Regression | 42.5 | 0.5 |
| PLEC-ANN | Regression | 35 | 0.411 |
| PLEC-DNN | Regression | 47.5 | 0.559 |
| MF-PLEC-RF | Regression | 52.5 | **0.617** |
| PLEC-RF | Classification | 32.5 | 0.382 |
| PLEC-XGB | Classification | 12.5 | 0.147 |
| PLEC-SVM | Classification | 30.0 | 0.352 |
| PLEC-ANN | Classification | 42.5 | 0.5 |
| PLEC-DNN | Classification | 27.5 | 0.323 |
| MF-PLEC-ANN | Classification | 43.8 | **0.515** |

### Limitations of the Butina approach

Despite the strong performance of several models, two limitations of the Butina split were identified.

#### Training on a single cluster of actives

The largest cluster, containing 33 of the 73 actives (45%), was used as the training set, so the models learned from actives that are structurally very similar to each other and had no exposure to other chemotypes. Training and test actives were also not fully separated: 35% of the test actives had a maximum Tanimoto similarity above 0.6 to at least one training active (median 0.50). Training on a single cluster limits the ability of the models to generalise to other regions of chemical space, and the test set is therefore a narrower assessment of generalisation than intended.

#### Hyperparameters were optimised on the training set

As the training set was small (119 compounds), no separate validation set was held out, and hyperparameters were optimised on the same data used to train the models. This provides no independent estimate of generalisation during tuning and increases the risk of overfitting to the training data, particularly for the more flexible models. Hyperparameter optimisation with a held-out validation set is the standard and recommended practice, and it motivated the use of cross-validation for hyperparameter optimisation in the UMAP-based approach.

#### Implications for the UMAP approach

These results motivated four changes in the UMAP-based approach. First, regression outperformed classification for most algorithms and gave the best model overall, so only regression was used. Second, the ANN and DNN models were the least stable and the training set was small, so only RF, XGB and SVM were used. Third, MF-PLEC gave the most consistent improvement over PLEC, whereas Smina terms and physicochemical descriptors mostly lowered performance, so PLEC and MF-PLEC were retained as the main PLEC-based schemes. Fourth, the two limitations above motivated a split with several training-test partitions, each testing on a held-out cluster of actives, and hyperparameter optimisation by stratified cross-validation on held-out portions of the training data, instead of on the training data itself.

### UMAP approach

The UMAP clustering split was designed to address the two limitations of the Butina split identified above.

#### Training on several chemotypes

In the Butina split, the models were trained on a single cluster of 33 structurally similar actives. In the UMAP splits, the training set of each split contains the actives of three of the four clusters (between 48 and 66 actives), so the models are exposed to several chemotypes. Each cluster is held out once, so every active is used as a test active exactly once and the conclusions do not rest on a single, arbitrary training-test partition. The structural separation between training and test actives varied between the splits. The median maximum Tanimoto similarity of test actives to training actives was 0.62, 0.16, 0.54 and 0.52 for clusters 0 to 3, and 58%, 0%, 28% and 0% of them exceeded 0.6, respectively, compared with a median of 0.50 and 35% for the Butina split. The UMAP splits therefore provide test sets spanning a range of structural separation, from the highly dissimilar actives of cluster 1 to those of cluster 0, which are more similar to their training set than the Butina test actives.

#### Hyperparameter optimisation on held-out data

In the Butina split, hyperparameters were optimised on the data used to train the models. In the UMAP splits, they were optimised by stratified 3-fold cross-validation within each training set, so that each candidate configuration was scored on compounds not used to fit it. The held-out cluster remained entirely unseen during both training and tuning, and is used only for evaluation.

The UMAP experiments were conducted in stages, each motivated by the results of the previous one: first the models were trained without PU decoys, then PU decoys were added to the training set, then additional featurisation schemes were evaluated, and finally RFE feature selection was applied to the best-performing featurisation scheme.

### Models trained without PU decoys

In the first set of UMAP experiments, the RF, XGB and SVM regression models were trained on the actives of three clusters and the 86 experimental inactives, without PU decoys. Overall, enrichment was low (Figure 7). The SVM was the best algorithm on clusters 0 and 3 (NEF1% of 0.363 and 0.666 on the matched-decoys test sets with PLEC) and the only model with any enrichment on cluster 2 (up to 0.166). On cluster 1, whose actives were the most dissimilar to the training actives, only the XGB model with PLEC achieved NEF1% above 0 (0.058 and 0.142 on the all-decoys and matched-decoys test sets, respectively). Values were generally higher on the matched-decoys test sets than on the all-decoys test sets. As the matched-decoys sets are much smaller, their top 1% contains few compounds and the values are coarse, particularly for cluster 3 (only 7 actives).

**Figure 7:**
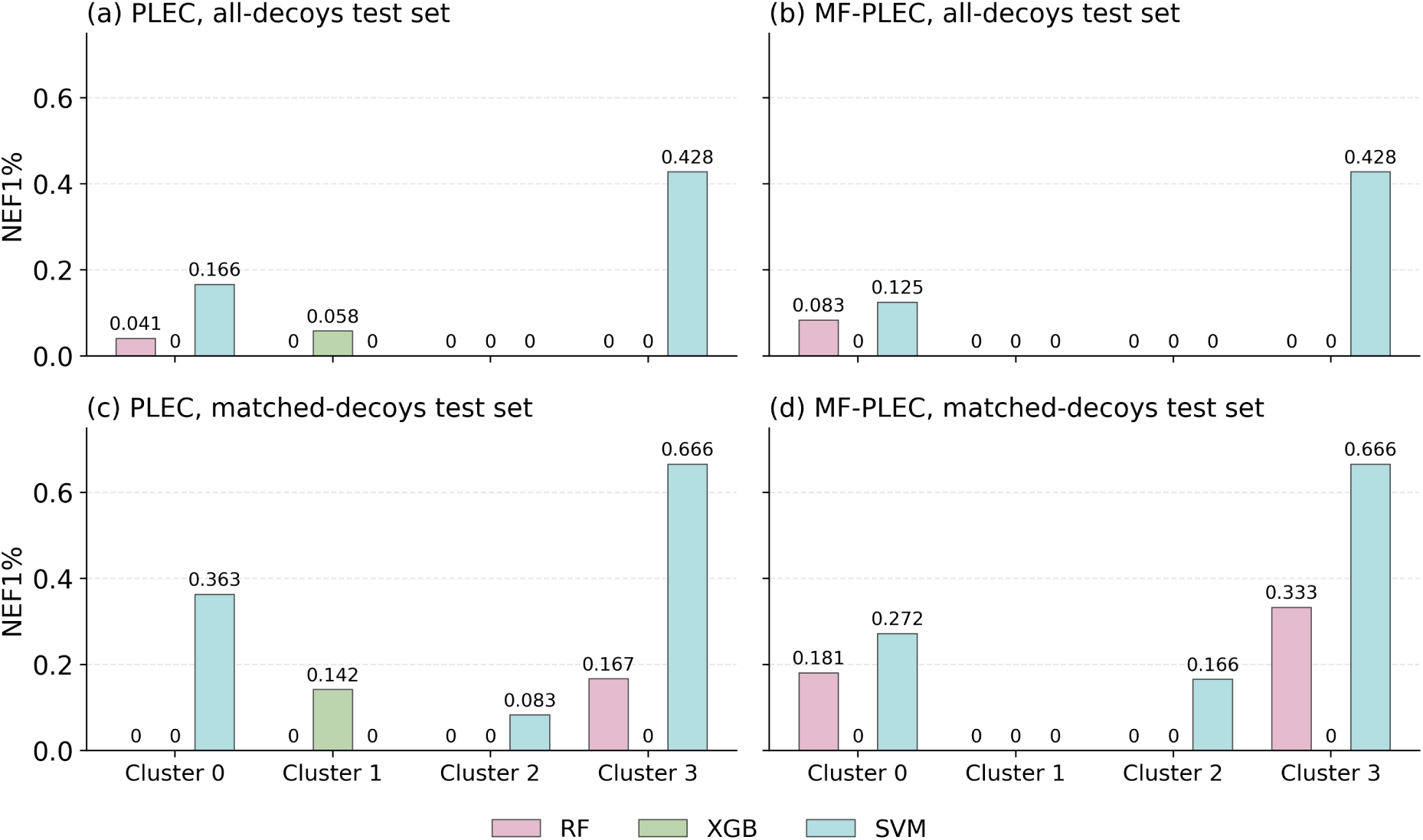
Median NEF1% over 10 runs for the *Pf*DHODH-specific RF, XGB and SVM regression models trained without PU decoys, for each held-out cluster of the UMAP clustering split (clusters 0 to 3). Models were trained with PLEC (a, c) or MF-PLEC (b, d) features and evaluated on the all-decoys test sets (a, b) and the matched-decoys test sets (c, d). A value of 0 indicates no enrichment in the top 1%.

Adding Morgan fingerprints (MF-PLEC) gave no consistent improvement. It increased the RF values on clusters 0 and 3 (matched-decoys test sets: 0 to 0.181 and 0.167 to 0.333) and the SVM value on cluster 2 (0.083 to 0.166), but lowered the SVM value on cluster 0 (0.363 to 0.272) and removed the XGB enrichment on cluster 1. The SVM was the best algorithm with either featurisation, and PLEC alone was often as good as MF-PLEC. Performance did not simply track the structural separation between training and test actives: cluster 1, the most dissimilar (median maximum similarity 0.16), gave the poorest results, but cluster 0, the most similar (0.62), was not the best.

Enrichment was near zero for clusters 1 and 2 with every model, meaning that decoys were ranked above most of the held-out actives at the top of the ranking. This could reflect decoys being scored too highly or the held-out actives being scored too low, so the predicted potencies of the held-out actives and the decoys were compared for each cluster (Figure 8). Actives were predicted to be more potent than decoys on average in all four clusters, so the models retained signal separating the two classes. The separation was clearest for cluster 3 (7 actives) and reasonable for cluster 0. Clusters 1 and 2 were the worst predicted, with many outliers among the predicted potencies of the decoys, some of which exceeded the potencies predicted for the actives. As early enrichment is determined by the highest-scored compounds, these decoy outliers explain the low NEF1% despite the higher average potency predicted for the actives. The problem therefore lay mainly with the decoys being poorly predicted, not with the actives being scored low. This suggested that the experimental inactives (86 compounds) used for training cover the inactive chemical space too narrowly, and that adding PU decoys to the training set to increase its chemical diversity might improve the separation of the two classes, which motivated the experiments described next.

**Figure 8:**
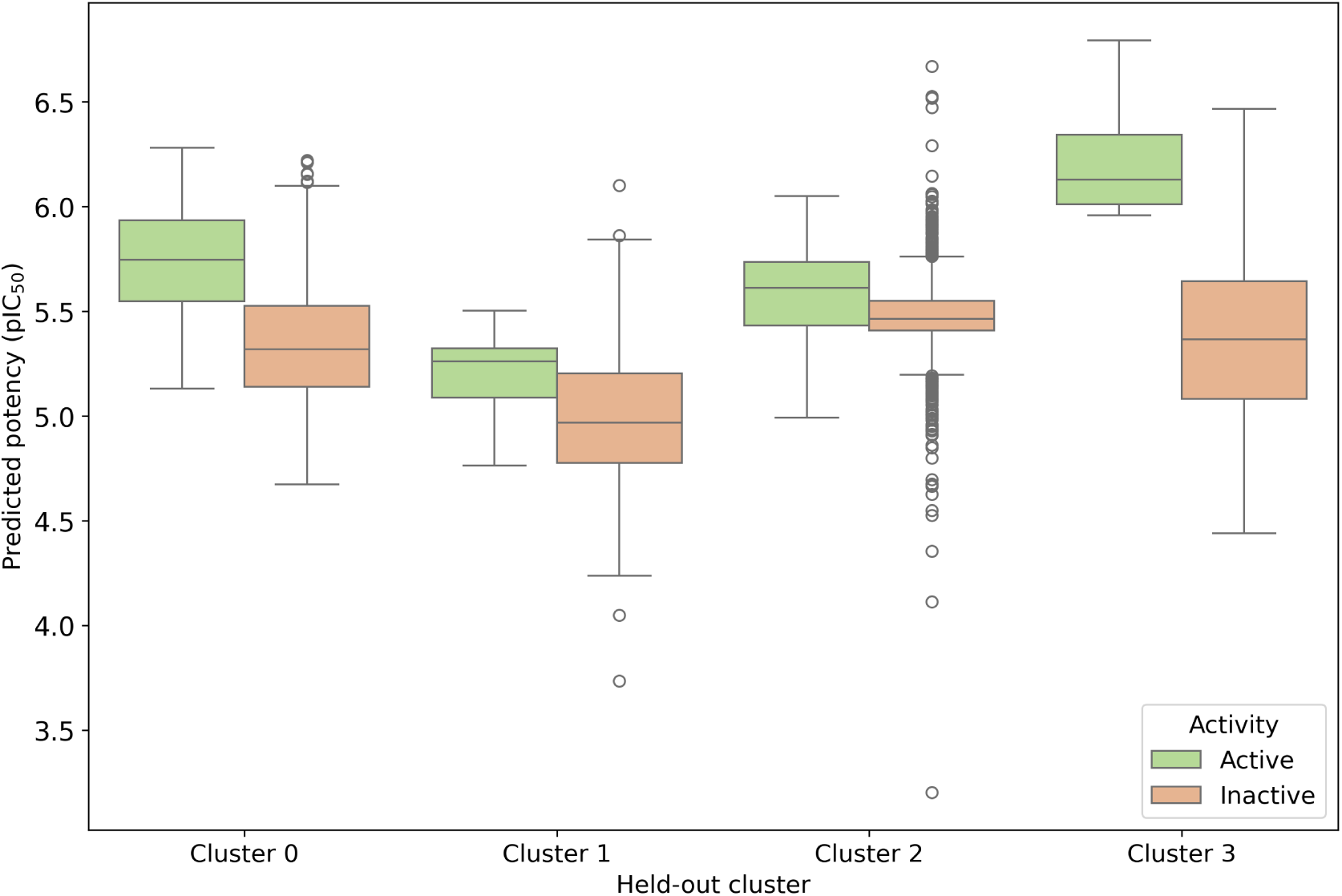
Predicted potency (pIC_50_) of the held-out actives and of their matched PM decoys for each cluster of the UMAP clustering split, shown for the matched-decoys test sets. Predictions were made with PLEC-SVM, the best-performing model overall, trained without PU decoys. Boxes show the interquartile range, lines the median, whiskers 1.5 times the interquartile range, and circles outliers.

### Effect of adding PU decoys to the training set

The poor discrimination of decoys observed above suggested that the experimental inactives alone describe the inactive chemical space too narrowly. To test this, the 1500 PU decoys were added to the training set of each split, and the models were retrained and re-evaluated on the same test sets.

The addition of PU decoys improved the performance of most models on clusters 0, 2 and 3, with both featurisation schemes and on both test sets (Figure 9). The largest gains were for clusters 2 and 3, where the RF and SVM models, which had NEF1% at or near 0 on cluster 2 without PU decoys, reached values of 0.88 to 1.0. The improvement was also clear for cluster 0 (for example, RF with PLEC from 0.041 to 0.333 on the all-decoys test set). These values should be read with caution for cluster 3, which has only seven actives, so that the top 1% of the matched-decoys test set contains about three compounds and NEF1% can take only a few discrete values. RF and SVM performed best, whereas XGB improved less and, with MF-PLEC, showed no improvement except on cluster 2 (about 0.25). Cluster 1 was the exception: NEF1% remained at or close to 0 for all models (at most 0.142 on the matched-decoys test set), despite the added decoys. As the actives of this cluster were the most dissimilar to the training actives, this suggests that the limitation for this cluster lies mainly in the representation of the actives rather than the inactives, which cannot be remedied by adding decoys.

**Figure 9:**
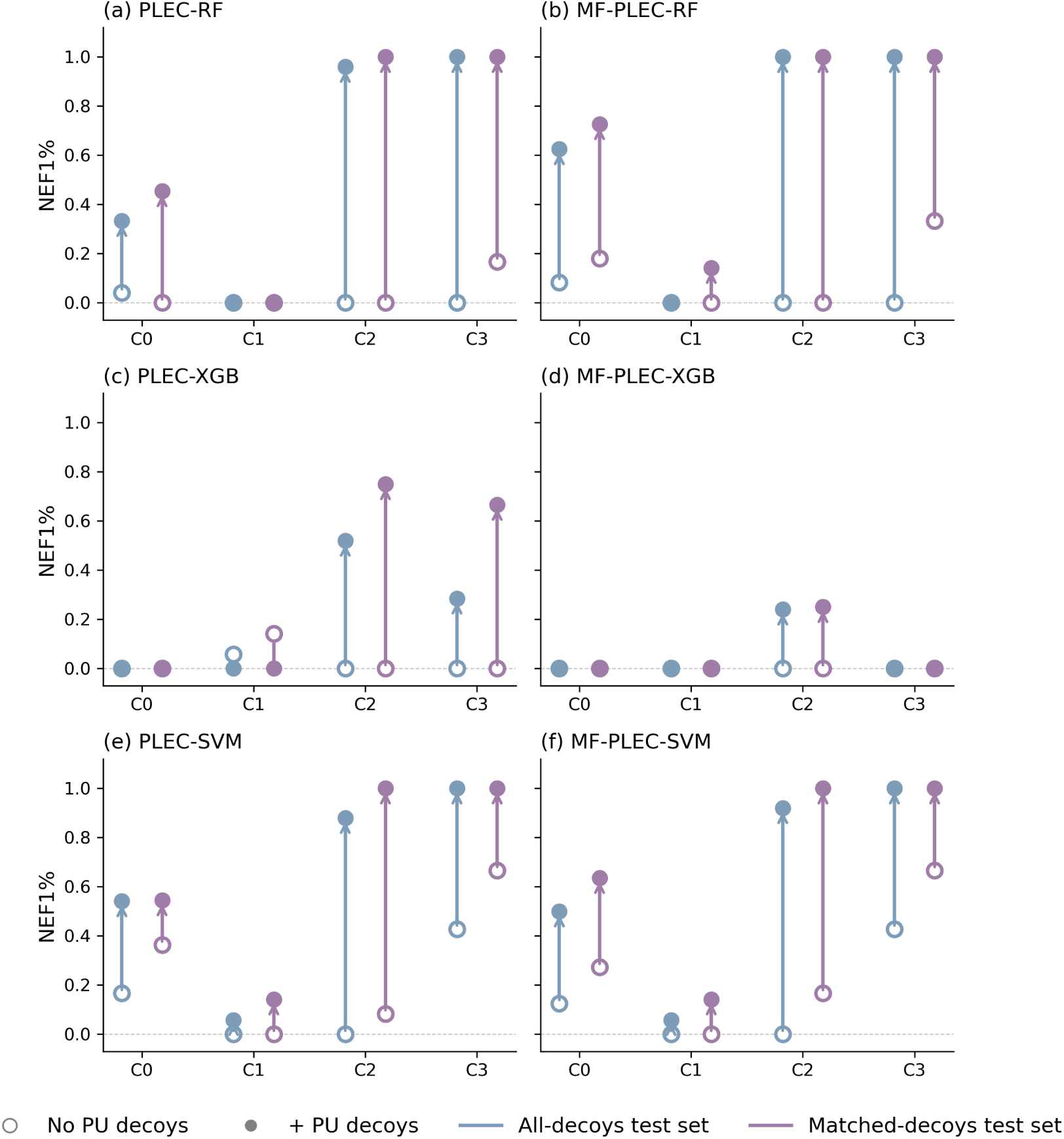
Effect of adding PU decoys to the training set on the median NEF1% over 10 runs for the *Pf*DHODH-specific RF (a, b), XGB (c, d) and SVM (e, f) regression models, with PLEC (a, c, e) and MF-PLEC (b, d, f) features, for each held-out cluster (C0 to C3) of the UMAP clustering split, on the all-decoys (blue) and matched-decoys (purple) test sets. Open markers show models trained without PU decoys and filled markers models trained with PU decoys, and arrows show the change in performance. Where the two markers coincide, only the filled marker is visible.

### Additional featurisation techniques

After establishing that PU decoys improve performance, two further protein-ligand interaction featurisation schemes, weighted and multi-shelled ECIF [33], were evaluated against PLEC (Figure 10). Performance was assessed on each cluster individually and then averaged across the four clusters.

**Figure 10:**
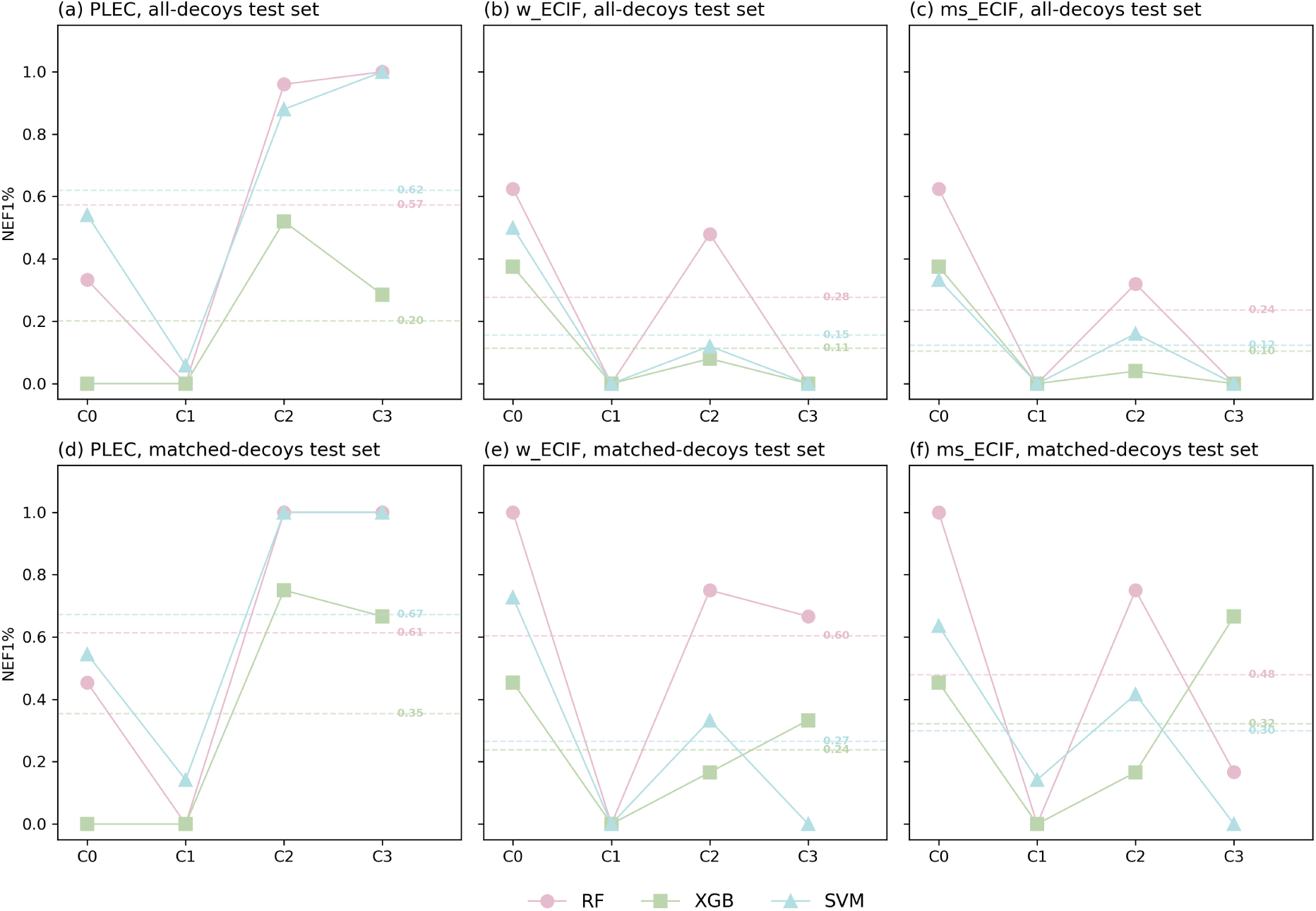
Median NEF1% over 10 runs for the *Pf*DHODH-specific RF, XGB and SVM regression models trained with PU decoys, using PLEC (a, d) weighted ECIF - w ECIF (b, e) and multi-shelled ECIF - ms ECIF (c, f) features, for each held-out cluster (C0 to C3) of the UMAP clustering split. Results are shown for the all-decoys (a-c) and matched-decoys (d-f) test sets. Lines join the clusters for visual clarity and dashed horizontal lines mark the mean NEF1% across the four clusters for each algorithm.

The ECIF schemes improved performance on cluster 0 for RF and XGB on both test sets, and for SVM on the matched-decoys test set, but generally performed worse than PLEC on clusters 2 and 3, and all models remained at or near 0 on cluster 1. Averaged across the four clusters, PLEC-SVM remained the best model (NEF1% of 0.62 on the all-decoys and 0.67 on the matched-decoys test set), followed by PLEC-RF (0.57 and 0.61). The ECIF models averaged between 0.11 and 0.28 on the all-decoys test set and between 0.24 and 0.60 on the matched-decoys one, where weighted ECIF with RF (0.60) came close to PLEC-RF. As PLEC were found to be the best-performing protein-ligand interaction featurisation scheme, additional investigation was carried out to test whether feature selection combined with PLEC can be useful.

### Feature selection with PLEC

To assess whether feature selection can benefit the models, RFE was applied to the PLEC features, the best-performing protein-ligand interaction featurisation scheme in the previous stage (Figure 11). Its effect was small and inconsistent, positive for some combinations of algorithm and held-out cluster and negative for others. The clearest improvement was on cluster 2 of the all-decoys test set, where RFE increased the NEF1% of all three algorithms (by 0.04 for RF and SVM and by 0.12 for XGB). On cluster 0 of the same test set, the effect differed between algorithms, with decreases for RF (-0.08) and SVM (-0.04) and an increase for XGB (+0.04). On the matched-decoys test set, RFE changed the NEF1% of XGB by +0.09 and of SVM by -0.09 on cluster 0, and of XGB by -0.08 on cluster 2. On cluster 1, the only change was a decrease for SVM on the all-decoys test set (-0.06), and there was no effect on cluster 3 on either test set. Averaged across the four clusters, the change in NEF1% (ΔNEF1%) was at most 0.04 in magnitude for any algorithm on either test set. Reducing the PLEC fingerprint from 4092 to 500 features therefore largely preserved performance on average, which suggests that many of the PLEC features are redundant for this task.

**Figure 11:**
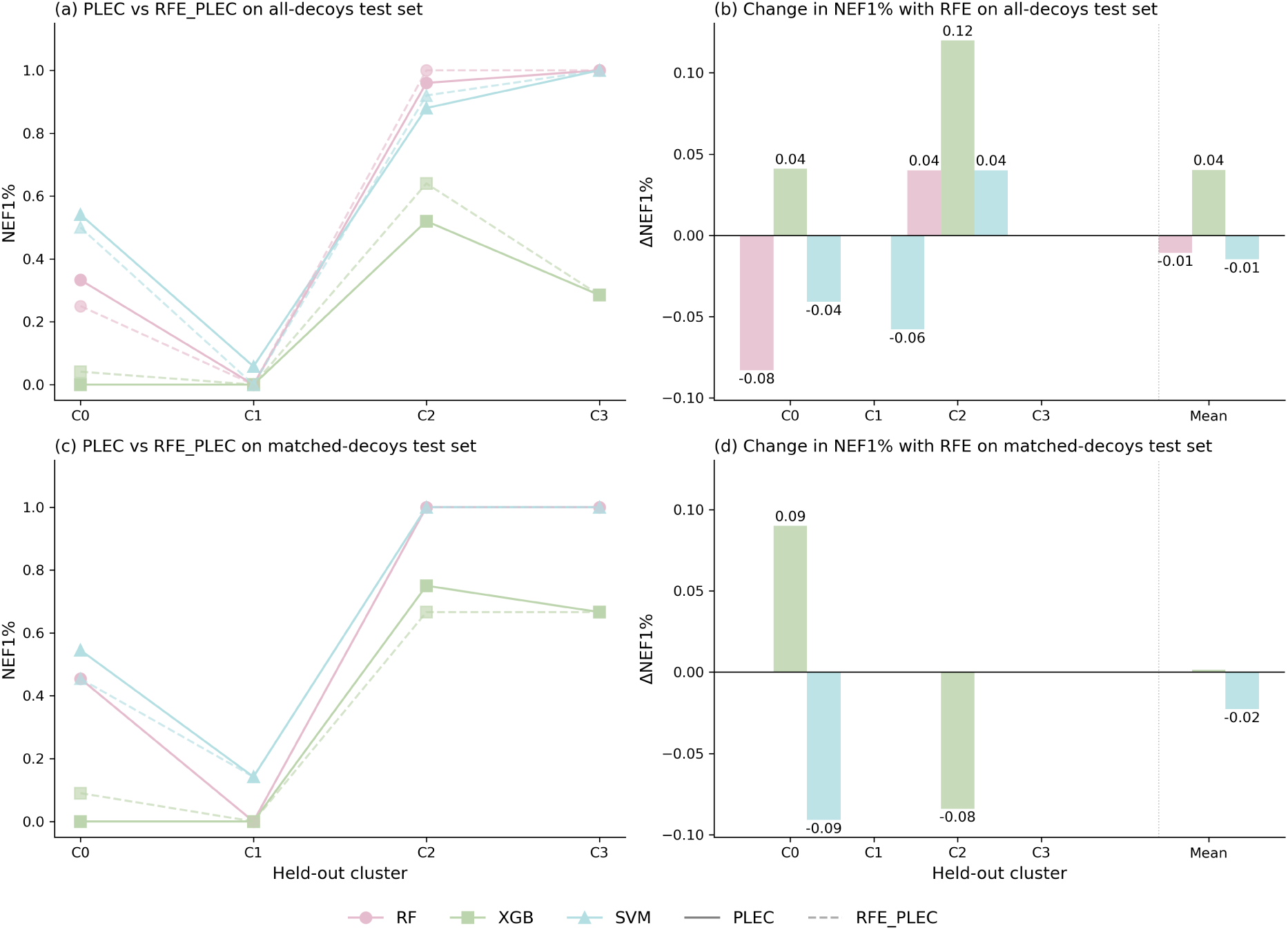
Effect of RFE on PLEC-based models trained with PU decoys, for each held-out cluster (C0 to C3) of the UMAP clustering split. (a, c) median NEF1% over 10 runs for RF, XGB and SVM with all 4092 PLEC features (solid lines) and with the 500 features retained by RFE - RFE PLEC (dashed lines), on the all-decoys (a) and matched-decoys (c) test sets. (b, d) change in NEF1% upon RFE (ΔNEF1%) for each algorithm and cluster, and the mean across the four clusters on the all-decoys test set (b) and matched-decoys test set (d). Lines join the clusters for visual clarity. Absent bars indicate no change.

### Comparison with reference SFs

The target-specific models were compared with the reference SFs using the NEF1% averaged across the four held-out clusters as the overall performance measure. Smina, CNNscore and CNNaffinity were included as reference SFs. The *Pf*DHODH-specific EHIGN was not carried forward to this comparison, as it requires training and did not outperform Smina and was outperformed by 8 out of 10 PLEC-based target-specific ML SFs in the Butina experiments.

On the all-decoys test set (Figure 12a), the best-performing model was MF-PLEC-RF, with an average NEF1% of 0.656. Among the models using PLEC alone, SVM outperformed RF, both with the full feature set (PLEC-SVM, 0.620, compared with 0.573 for PLEC-RF) and after feature elimination (RFE_PLEC-SVM, 0.605, compared with 0.562). Adding Morgan fingerprints did not improve the SVM (MF-PLEC-SVM, 0.620). Among the reference SFs, CNNscore and CNNaffinity performed poorly (0.032 and 0.051, respectively), whereas Smina achieved a moderate 0.362, higher than all XGB-based models and all models trained with multi-shelled or weighted ECIF features.

**Figure 12:**
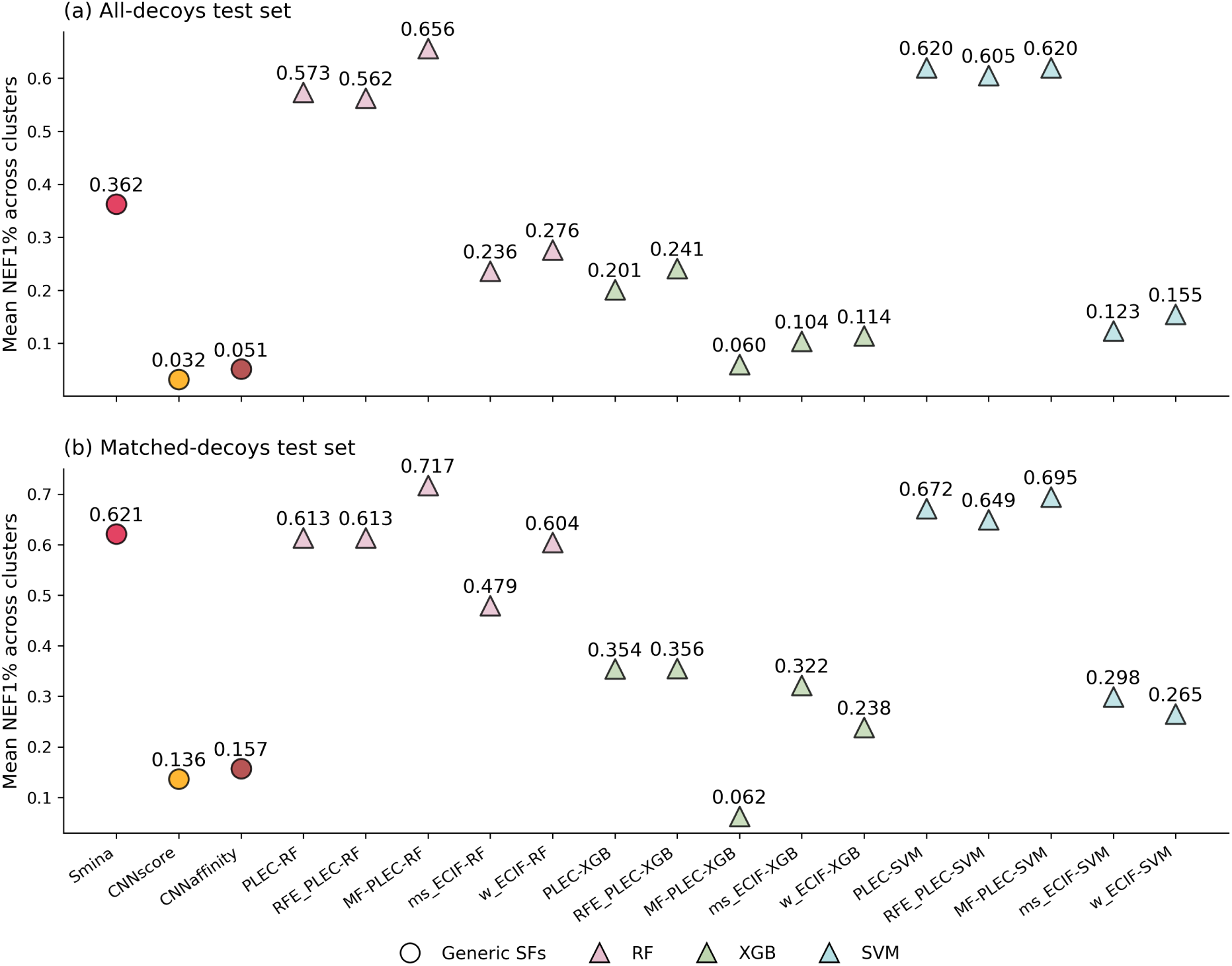
Mean NEF1% across the four held-out clusters of the UMAP clustering split for the generic SFs (Smina, CNNscore and CNNaffinity) and the *Pf*DHODH-specific RF, XGB and SVM regression models trained with PU decoys, on (a) the all-decoys and (b) the matched-decoys test sets. Target-specific models are shown for PLEC, PLEC with RFE (RFE_PLEC), MF-PLEC, multi-shelled ECIF (ms_ECIF) and weighted ECIF (w_ECIF). For the target-specific models, the per-cluster values used to calculate the mean are medians over 10 runs.

On the matched-decoys test set (Figure 12b), MF-PLEC-RF was again the best target-specific model (0.717), followed by MF-PLEC-SVM (0.695) and PLEC-SVM (0.672). All methods scored higher on this test set than on the all-decoys test set, which is expected given its smaller size and higher fraction of actives. Smina performed very well here (0.621) and was outperformed only by the MF-PLEC-RF and the SVM models with PLEC-based features (PLEC, RFE_PLEC and MF-PLEC). All other target-specific models scored lower than Smina, including PLEC-RF and RFE_PLEC-RF (0.613 each).

## Conclusions

This work presented, to our knowledge, the first target-specific ML SFs developed for *Pf*DHODH. In the Butina experiments, most PLEC-based models outperformed Smina, CNNaffinity, CNNscore and the *Pf*DHODH-specific EHIGN, with the best model (the RF regressor with MF-PLEC) reaching a NEF1% of 0.617, compared with 0.323 for Smina. Smina was the best-performing generic SF on this target in both the Butina and the UMAP experiments, with CNNscore and CNNaffinity falling well behind, and it was therefore used as the main comparator. Regression models generally outperformed their classification counterparts, and the ANN, DNN and EHIGN models, which are more data-hungry, were unstable or did not outperform Smina with the small training set available. The Butina split also exposed two limitations of the evaluation: the models were trained on a single cluster of structurally similar actives, and hyperparameters were optimised on the training data itself. These motivated the UMAP-based approach.

The UMAP splits trained the models on several chemotypes and held out each cluster of actives in turn, so performance could be assessed across actives of varying similarity to the training set, with hyperparameters optimised by cross-validation. Performance varied strongly between clusters. Without PU decoys, enrichment was low and close to zero for clusters 1 and 2, because decoys were poorly predicted. Adding PU decoys to the training set increased the chemical diversity of the inactive class and substantially improved enrichment for clusters 0, 2 and 3, which shows their value for targets with few experimentally confirmed inactives. Cluster 1, the most dissimilar to the training actives, remained at or near zero, indicating that the models generalise poorly to chemotypes that are not represented in the training data.

PLEC remained the most competitive protein-ligand interaction featurisation, with the weighted and multi-shelled ECIF variants performing worse on average, although they improved some models on cluster 0. The best model overall was the RF regressor with MF-PLEC (average NEF1% of 0.656 and 0.717 on the all-decoys and matched-decoys test sets), followed closely by the PLEC-SVM (0.620 and 0.672). Notably, the MF-PLEC-RF configuration was also the best-performing under the initial Butina split (NEF1% = 0.617), suggesting that the best method does not depend on the splitting strategy. RFE reduced the PLEC features from 4092 to 500 with a small although inconsistent effect on performance, suggesting that many of the PLEC features are redundant. The best target-specific models exceeded Smina on both test sets, but the margin was much smaller on the matched-decoys test set, where Smina (0.621) outperformed most of the target-specific models.

Several limitations need to be acknowledged. Firstly, the dataset was small (73 actives and 86 experimental inactives), and the held-out clusters were smaller still, so the metrics are coarse, especially for cluster 3 (seven actives), where a single compound changes NEF1% considerably. Secondly, the experiments were conducted sequentially, with each stage motivated by the previous results and with several choices informed by test set performance, so not every combination was evaluated and the comparisons between stages are exploratory. Thirdly, a single crystal structure (4RX0) was used as the docking template. Fourthly, PU decoys were assumed to be inactive without experimental confirmation. Finally, the models have not been validated prospectively, so it is not yet known whether the retrospective enrichment would translate into experimentally confirmed hits.

A further limitation concerns selectivity. Since inhibition of human DHODH is undesirable for an antimalarial drug, incorporating selectivity, for example by training models on human DHODH data and prioritising compounds predicted to be active against the parasite enzyme only, is a natural extension of this work.

Overall, it has been shown that target-specific ML SFs can be developed for a target with scarce bioactivity data, provided that the evaluation is designed with care. The insights that PU decoys help when experimental inactives are limited, that regression and simpler algorithms are preferable with small training sets, and that the splitting strategy strongly affects the conclusions that can be drawn, are likely to be useful for developing such SFs for other targets.

